# A multimodal imaging and analysis pipeline for creating a cellular census of the human cerebral cortex

**DOI:** 10.1101/2021.10.20.464979

**Authors:** Irene Costantini, Leah Morgan, Jiarui Yang, Yael Balbastre, Divya Varadarajan, Luca Pesce, Marina Scardigli, Giacomo Mazzamuto, Vladislav Gavryusev, Filippo Maria Castelli, Matteo Roffilli, Ludovico Silvestri, Jessie Laffey, Sophia Raia, Merina Varghese, Bridget Wicinski, Shuaibin Chang, Anderson Chen I-Chun, Hui Wang, Devani Cordero, Matthew Vera, Jackson Nolan, Kim Nestor, Jocelyn Mora, Juan Eugenio Iglesias, Erendira Garcia Pallares, Kathryn Evancic, Jean Augustinack, Morgan Fogarty, Adrian V. Dalca, Matthew Frosch, Caroline Magnain, Robert Frost, Andre van der Kouwe, Shih-Chi Chen, David A. Boas, Francesco Saverio Pavone, Bruce Fischl, Patrick R. Hof

## Abstract

Cells are not uniformly distributed in the human cerebral cortex. Rather, they are arranged in a regional and laminar fashion that span a range of scales. Here we demonstrate an innovative imaging and analysis pipeline to construct a reliable cell census across the human cerebral cortex. Magnetic resonance imaging (MRI) is used to establish a macroscopic reference coordinate system of laminar and cytoarchitectural boundaries. Cell counting is obtained with both traditional immunohistochemistry, to provide a stereological gold-standard, and with a custom-made inverted confocal light-sheet fluorescence microscope (LSFM) for 3D imaging at cellular resolution. Finally, mesoscale optical coherence tomography (OCT) enables the registration of the distorted histological cell typing obtained with LSFM to the MRI-based atlas coordinate system.

## 1 Introduction

The human brain is a complex organ organized across a range of spatial scales. To understand its properties and functionality, it is critical to produce a comprehensive characterization of the neuronal cell types that compose it and to visualize their distributions through the whole volume^1,2^. Although significant technological advances ^3–7^ have been made to obtain complete cell census in animal models such as mouse and marmoset monkey^8–16^, no current imaging technology can directly visualize the defining microscopic features of the human brain without significant distortion. Indeed, available cytoarchitectural parcellations of the human brain^17^ are limited by unavoidable distortions introduced by slice-to-slice sectioning, clearing, staining and mounting steps involved in current histochemistry protocols. This processing results in artifacts that prevent accurate visualization of the tissue’s morphomolecular properties such as individual cells across different regions or laminar and cytoarchitectural boundaries that form the natural coordinate system for a cell census of the human brain.

Important steps have been made towards building 3D models of the human brain with cellular resolution through the use of magnetic resonance imaging (MRI)^18–30^, standard histology^31–38^ and polarized light imaging (PLI^39^). For example, the Big Brain project^17^ required 5 years and 1,000 person-hours to obtain a comprehensive dataset of one human brain with a nominal resolution of 20 μm. In parallel, data analysis and atlassing methods have been proposed to manage the the very large datasets generated during reconstructions and mapping of the volumes to standardized templates^6,40–42^. While these technologies represent remarkable advances, they still do not produce the undistorted 3D images of the cytoarchitecture of the human brain that are needed to build accurate models with nuclear and laminar resolution, a critical component of any cellular atlas.

Here, we present an innovative imaging and analysis pipeline that combines a number of imaging techniques to overcome the inherent limitations of each individual methodology. We propose to bridge microscopic volumetric histological imaging and macroscopic MRI using mesoscopic optical coherence tomography (OCT) to enable registration of the distorted histological volumes containing cell typing to an MRI-based atlas coordinate system. OCT is an optical technique that measures the backscattered light of the sample to provide high resolution cross-sectional imaging and 3D reconstruction up to several hundred micrometers in depth in fixed *ex vivo* biological tissues, in a contact-free and non-invasive manner (no staining required)^43^. Serial sectioning OCT can be accurately registered to the whole-hemisphere MRI of the brain as imaging is performed prior to slicing and thus virtually minimal deformation is introduced. For 3D histological analysis with even higher sub-cellular resolution, confocal light-sheet fluorescence microscopy (LSFM)^44^ is employed to image the large volumes of human brain tissue slabs obtained after OCT measurement. LSFM allows to obtain a fast optical sectioning of the sample by employing a specific configuration where the illumination axis is orthogonal to the acquisition axis^44–46^. LSFM is coupled with a dedicated tissue transformation technique to specifically label neuron subtypes and clear the tissue.

Finally, a stereological assessment is performed on the LSFM reconstructions to obtain a cell type-specific quantitative census of the neurons.

This multimodal imaging infrastructure was developed to bridge the resolution gap between macroscopic and microscopic techniques, resulting in a platform that integrates cellular anatomical information within a whole-brain reference space. The pipeline represents a large multi-institutional collaborative effort across the Massachusetts General Hospital, Boston University, the European Laboratory for Non-Linear Spectroscopy, and the Icahn School of Medicine at Mount Sinai.

## 2 Results

### 2.1 Imaging and Analysis Pipeline Overview

In this work, we generated a cell census of the human cerebral cortex by implementing a multimodal imaging and analysis pipeline on Broca’s area (Brodmann’s areas 44/45), blocked from whole human hemispheres. The postmortem specimen included in this project was obtained from a subject who had no neurologic or psychiatric illnesses. **Figure 1** provides an overview of the imaging pipeline performed on the sample (left, from top to bottom) and the subsequent imaging and data analysis pipeline (left to right). The whole hemisphere is first imaged with MRI before Broca’s area is blocked and imaged with serial sectioning OCT. 3D histological fluorescence imaging on the acquired slices is then performed with LSFM. In parallel, gold-standard stereology is performed on additional tissue samples. A stereological evaluation is performed digitally on the LSFM data with a laminar level of resolution. An atlas of the data is obtained through non-linear registration of the three modalities; LSFM to OCT and OCT to MRI. Cell counts and manually labeled features are also registered along with the volumes on which they were generated, thereby mapping the results of our analysis to a within-subject coordinate system. All the data collected are made available on the DANDI platform^47^. In the following sections, details are provided on each step of this pipeline.

**Figure 1:**
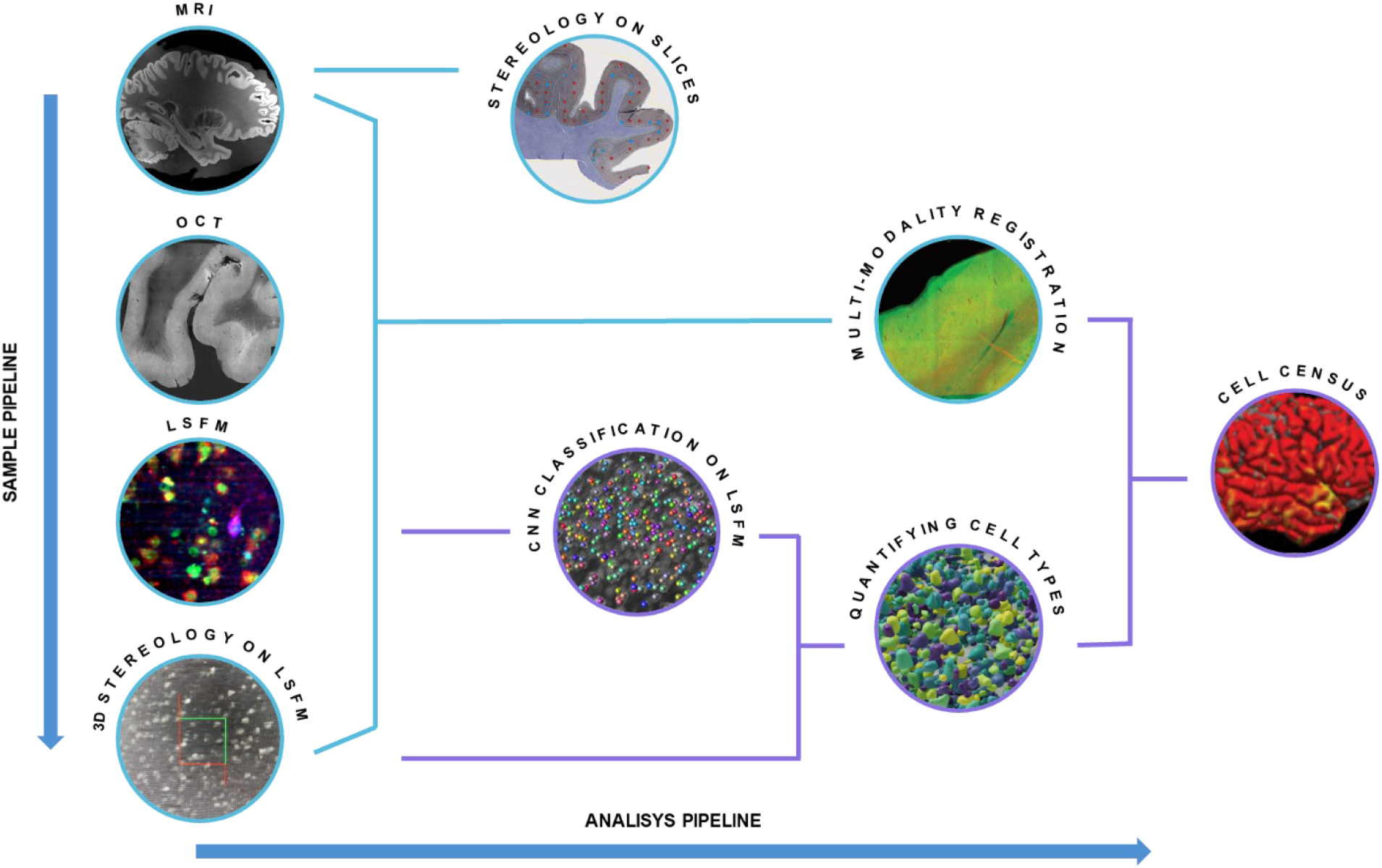
Imaging and analysis pipeline overview. From top to bottom sample pipeline: MRI, OCT, LSFM, and 3D stereology on LSFM images are performed on the same sample. Left to right analysis pipeline: thin-section stereology is performed on sections from separate specimens to obtain gold-standard counting; registration between MRI, OCT, LSFM and 3D sterology data is performed to align all the datasets back to the MRI coordinate system; cell counting with CNN classification on LSFM images will permit automatic counting of the stained neurons; the combination of 3D stereologic evaluation and CNN automatic counting will obtain a reliable quantification of cell types; multimodal registration between the images and the counting is needed to finally obtain the cell census of the neurons in a MRI-based atlas coordinate system. Schematic in cyan represents the steps presented in the current manuscript where a Broca’s Area block of tissue underwent to all the steps. Violet steps are future implementation of the pipeline.

### 2.2 Reference coordinate system establishment with MRI imaging

A reference coordinate system for the cellular atlas was established with *ex vivo* MRI of the whole hemisphere. Structural images with 150-μm isotropic resolution were acquired using a multi-flip angle, multi-echo fast low angle shot (FLASH) MRI sequence on a Siemens 7 T MRI scanner using a custom-built 31-channel head coil^21^. We performed diffusion imaging on a Siemens 3 T MRI scanner using a Siemens 32-channel head coil. The MRI data were processed to correct for geometric distortions due to *B*_0_ field inhomogeneities^48–50^, contrast variations due to *B*_1_ transmit field inhomogeneities and intensity bias due to *B*_1_ receive coil sensitivity variations^51^. **Figure 2** illustrates the 150-μm isotropic root mean squared (RMS) image for a flip angle of 20° calculated across the four echo times on the sample presented in this paper. **Figure 2** also demonstrates the improvement in vessel and laminar contrast quality attained by performing various artifact correction steps. **Figure 2a** zooms into the Broca’s area where a clear contrast between grey and white matter, and the transition from the infra-to supragranular layer of the cortex within the grey matter, is visible in these images. These laminar boundaries are critical for tabulating information about cell types and their distributions within the characteristic architectural infrastructure that defines the cortical sheet. **Figure 2b** shows a zoomed-in portion of the frontal lobe that is severely effected by *B*_0_ and *B*_1_ transmit distortions. The *B*_0_ inhomogeneity blurred the vessels and cortex in the RMS image due to misalignment between their locations in the different echo images. B1 transmit field variation reduced the overall vessel contrast. Vessels are important anatomical landmarks used to aid cross-modality registration and since the distortion corrected images demonstrate improved vessel sharpness and contrast, they aid in providing accurate landmarks essential for registration. **Figures 2c and 2d** show axial and sagittal slices before and after intensity bias correction. The bias-corrected images show improved laminar and white matter contrast. The artifact-corrected whole-hemisphere MRI data presented here thus provides a reliable laminar framework, improved quality of vessel landmarks, and a reference space to which downstream modalities can be registered.

**Figure 2:**
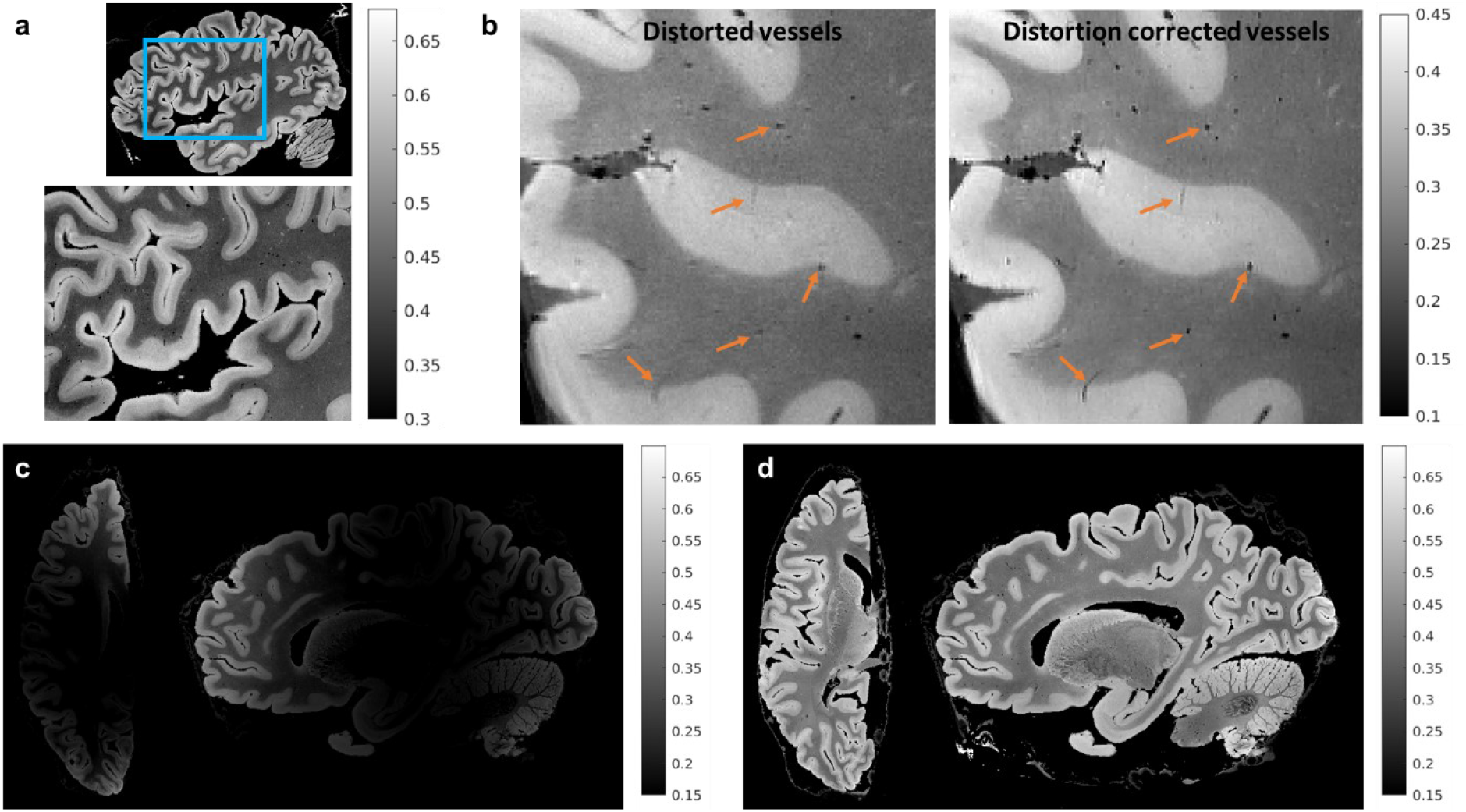
MRI for global reference. The figure shows results for 150 μm isotropic root mean squared (RMS) FLASH MR images with RMS calculated across four echo times (TE= 5.65, 11.95, 18.25 and 24.55 ms) for a flip anlge of 20°. a) A sagittal section that is zoomed into the Broca’s area showing infra- and supra-granular gray matter contrast. b) A zoomed-in sagittal frontal section of the same brain sample with vessels that are blurred and have reduced contrast. The blurring is due to B0 field inhomogeneity while the reduced contrast is due to B1 transmit field inhomogeneity. We also show vessels post-distortion correction that are well preserved and have high contrast, demonstrating the effectiveness of the correction methods. c) and d) show an axial and sagittal slice before and after B1 receive intensity bias correction. The intensities are visibly more uniform after bias removal.

### 2.3 OCT 3D reconstruction for tissue registration

The excised block (approximately 1.5×1.3×0.8 cm^3^) was imaged with a home-built serial sectioning OCT system at 5-μm isotropic mesoscale resolution that reveals cortical layers and cytoarchitectural boundaries^43^. To enhance the penetration of light deep inside the sample for OCT, we exploited a clearing procedure, based on the organic solvent 2,2’-thiodiethanol (TDE), to reach an imaging depth of up to 500 μm^52^. Sectioning the 500 μm-thick slices was performed using a custom-built vibratome that is capable of slicing sections up to 6 cm-wide^53^. By capturing the intrinsic back-scattering properties of the tissue, OCT elucidated features such as vasculature and cortical layer structure to be used as registration landmarks (**Figure 3** and **Supplementary Movie1**). Additionally, as OCT imaging precedes sectioning, tissue deformations from sectioning were almost completely eliminated.

**Figure 3:**
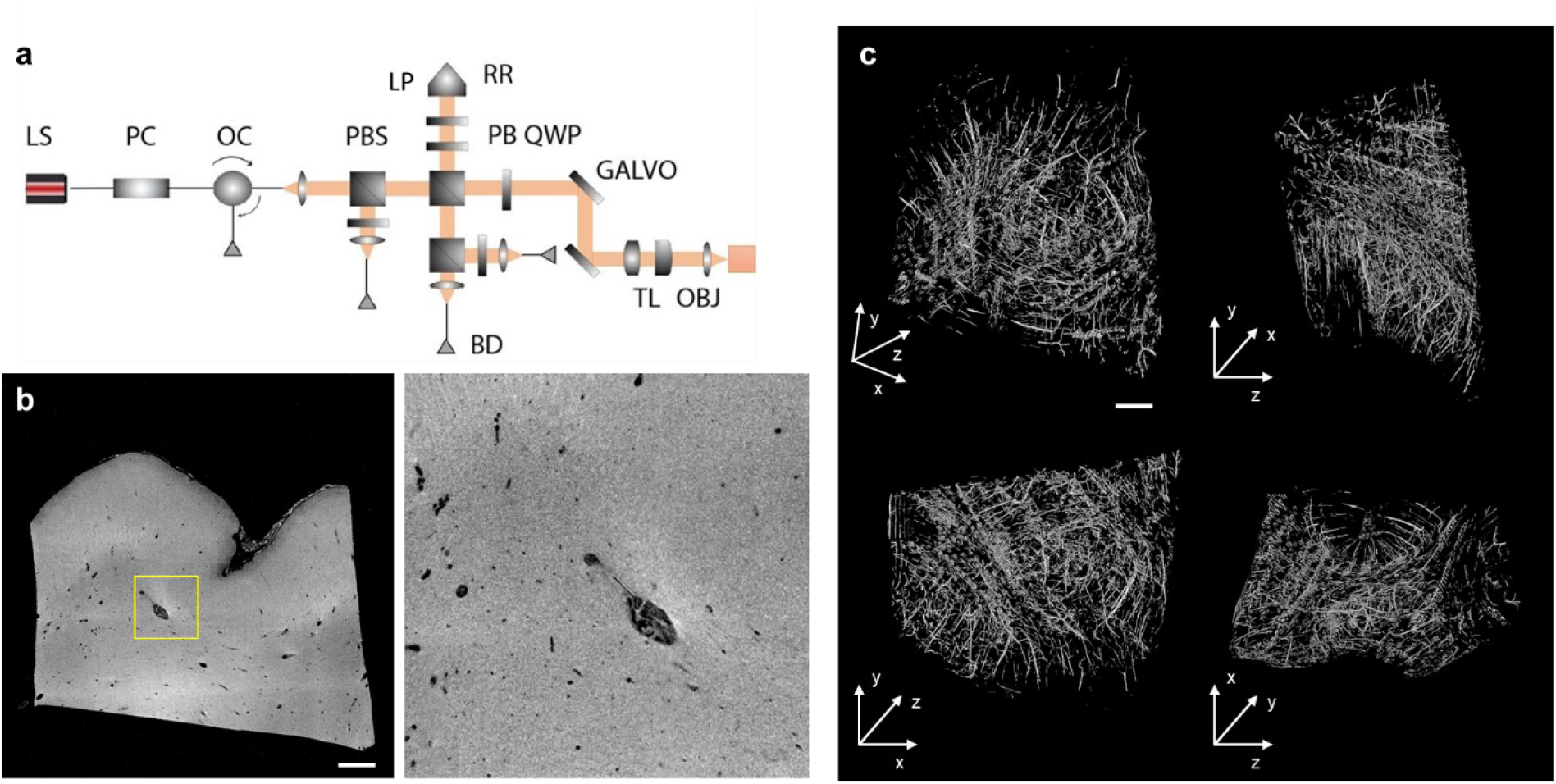
OCT analysis. a) Schematic of OCT. Annotations of components are LS: light source, PC: polarization controller, OC: optical circulator, PBS: polarizing beam splitter, PBS: polarization beamsplitter, RR: retroreflector, LP: linear polarizer,QWP: quarter wave plate, GALVO: galvo mirror, BD: balanced detector, TL: telescope, and OBJ: objective. b) An example XY slice of the OCT volume intensity (left) and zoom-in view of the highlighted window (right, dimension = 1.5 x 1.5 mm). Scale bar: 1 mm. c) 3D rendering and orthogonol views of the vessel segmentation of the OCT volume using the Frangi filtering method. Top left: 3D rendering view. Top right: YZ view. Bottom left: XY view. Bottom right: XZ view. Scale bar: 1 mm.

### 2.4 Molecular phenotyping reconstruction with LSFM

Fluorescence imaging of the 16 slices was obtained using a custom-built dual-view inverted confocal LSFM with a voxel-size resolution of 0.55 x 0.55 x 3.3 μm^3^ that results in a 3.3 μm^3^ isotropic resolution after post-processing (**Figure 4a**)^54^. Molecular specificity was achieved by combining LSFM with an advanced tissue transformation protocol called SHORT^55^. The protocol rendered the sample completely transparent to light by refractive index matching (**Figure 4b**) and allowed homogeneous co-labeling of large 3D volumes with different markers (**Figure 4c** and **Supplementary Figure 1**). To perform the cell census in the area 44/45, we used immunofluorescence to label specific neuronal populations. The use of neurochemical markers allows for the definition of region-specific staining patterns, and generally results in a high definition of cortical areas that complement traditional Nissl and myelin preparations. In this context, calcium-binding proteins have been shown to be reliable cellular markers for cytoarchitectural studies of the primate neocortex, in which they are present principally in distinct populations of inhibitory neurons that exhibit recognizable patterns of regional and laminar specialization^35,56–64^. We used an anti-neuronal nuclear antigen (NeuN) antibody to stain all neurons and an anti-calretinin (CR) antibody to identify a subpopulation of γ-aminobutyric acid-ergic (GABAergic) interneurons. To detect all cell nuclei we used an exogenous dye (propidium iodide, PI) obtaining a three-color costaining in the same tissue (**Figure 4d and Supplementary Movie 2**). Vessels were identified from autofluorescence signals generated by retained blood.

**Figure 4:**
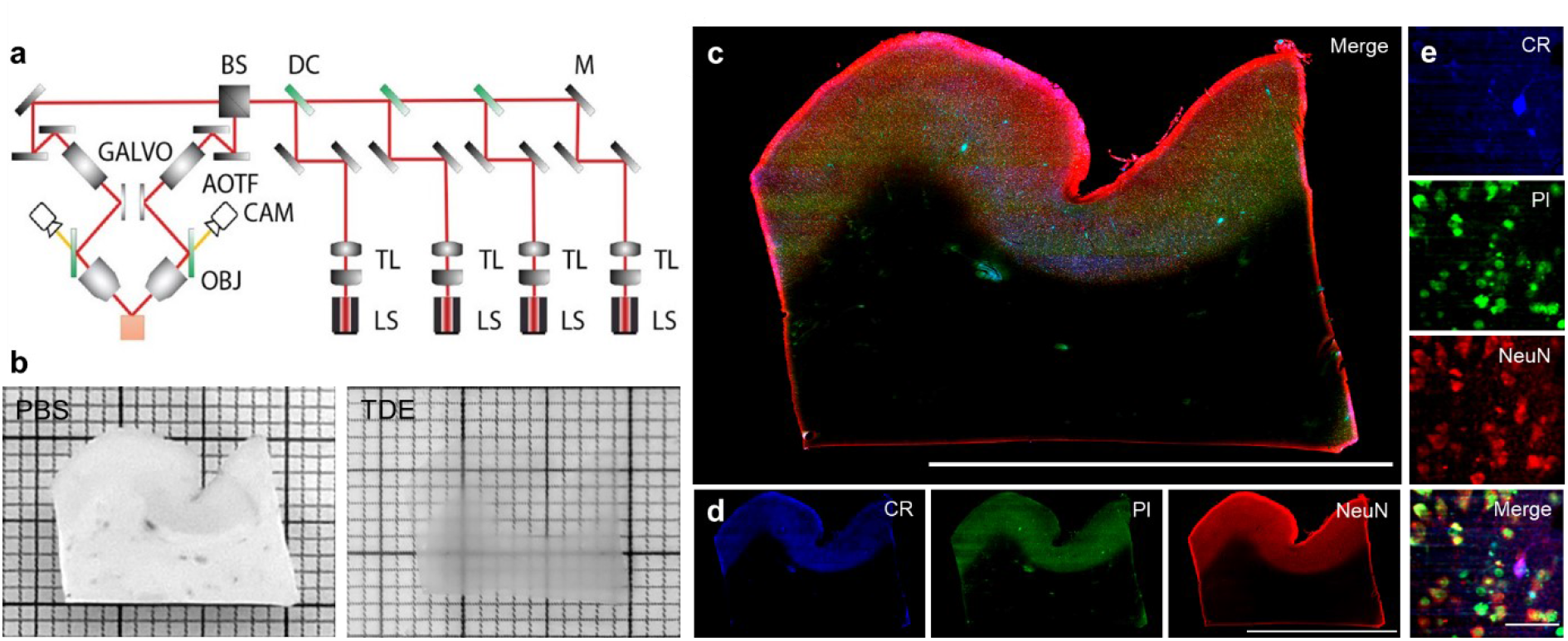
Multicolor imaging with LSFM. a) Schematic of LSFM apparatus. Annotations of components are LS: laser source, TL: telescope, M: mirror, DC: dichroic, BS: beam splitter, GALVO: galvo mirror, AOTF: acousto-optical tunable filters, CAM: camera, and OBJ: objective. b) A 500 μm-thick slice in PBS before (left) and after (right) TDE tissue clearing. c) A representative slice of a middle plane (≈200 μm depth) of a 500μm-thick slice stained with CR in blue (λ_exc_ =488), PI in green (λ_exc_ =561), and NeuN in red (λ_exc_ =638). Vessels are visible in the blue and green channels due to the presence of autofluorescence signals. d) Single channels of slice in c. Scale bars = 1 cm. e) High-resolution insets showing the different cellular markers used in the study: calretinin (CR), propidium iodide (PI), neuronal nuclear antigen (NeuN). Scale bar = 50 μm.

### 2.5 Stereological validation of 3D reconstruction

While LSFM enables the investigation of brain structures from the subcellular to the mesoscale, by recording different distributions of neuronal populations in large, cleared, specifically immunolabeled tissues with micrometric resolution and reasonably efficient acquisition times, no quantitative gold-standard validations of such population-level imaging data acquisitions currently exist. Hence we performed stereological gold-standard assessments on LSFM reconstructions of identified neuronal populations using an Optical Fractionator probe^65,66^ adapted to cleared materials (**Figure 5** and **Supplementary Figure 2**). We recorded on average 22,400 CR^+^ neurons in layer 3 across the slabs, 85,500 NeuN^+^ neurons in layer 3 and 62,700 NeuN^+^ in layer 5, with corresponding densities of 2787, 8120, and 6168 neurons/mm^3^, respectively (**Supplementary Table 1**).

**Figure 5:**
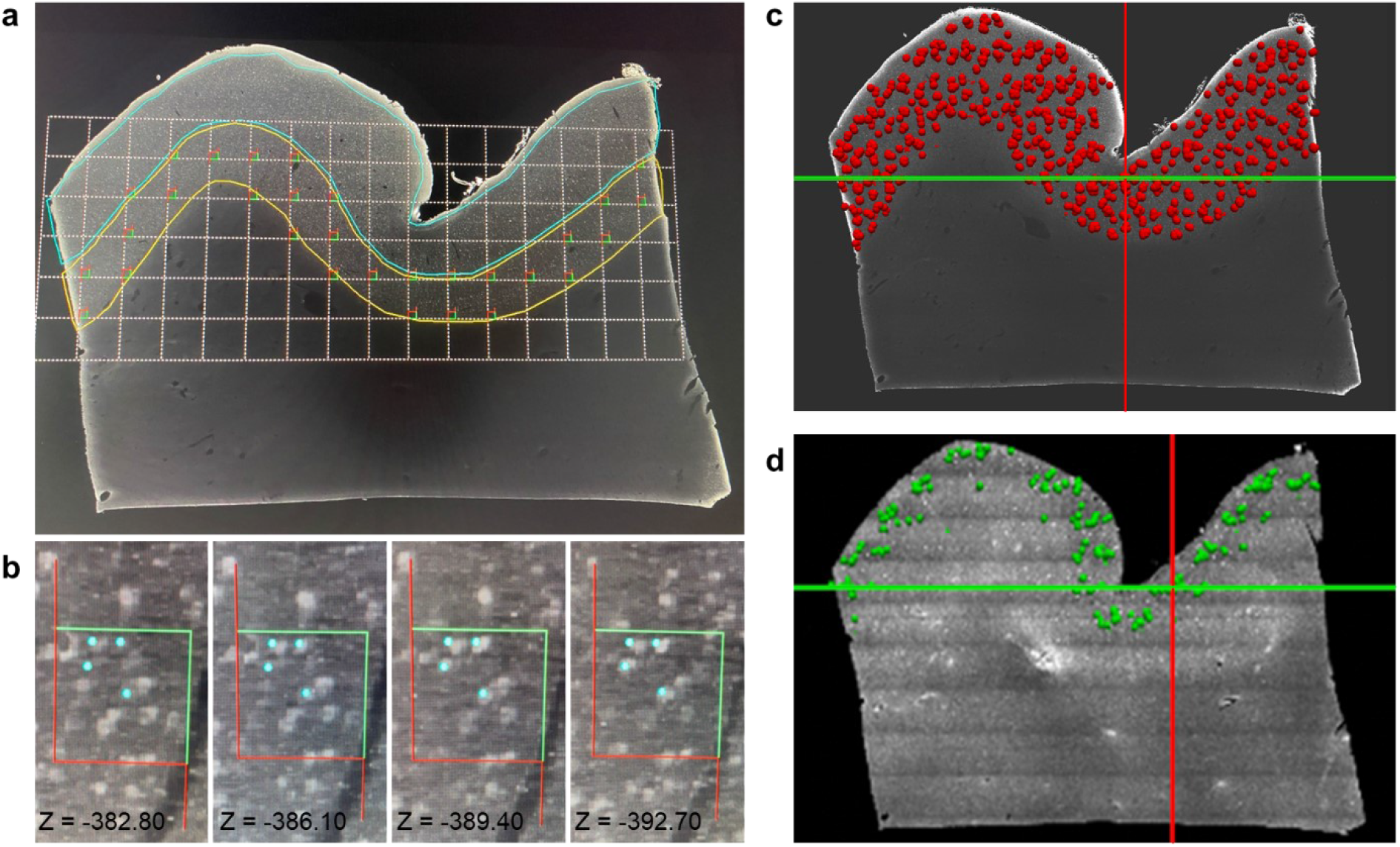
Stereological evaluation on LSFM reconstructions. a) A slab from the LSFM imaging dataset stained with the anti-NeuN antibody, displaying a systematic-random counting grid employed during stereologic analyses. The grid is placed over the infragranular layers (yellow outline) and optical disector frames are placed in all segments of the grid that are located within the region of interest. Panel b) shows different z-levels within a disector with blue markers indicating cells that have been sampled according to stereologic rules^38,39^. Panels c) and d) represent the registration of stereologic data onto the LSFM dataset for the entire slab. Each red dot in c) corresponds to NeuN^+^ neurons sampled through the slab within an optical disector depth during stereologic analysis and used to generate an estimate of the total population number. The green dots in d) represent CR^+^ neurons that were analyzed only in layers 2 and 3 as they are very sparse in the deep layers of the neocortex. Grid size on (a) = 800 μm, counting frame on (b) = 150 μm.

These LSFM data were validated against stereologic estimates from separate specimens prepared using traditional protocols and analyzed under brightfield conditions (see supplementary materials). This provides the ability to compare manual estimates to automatic segmentations obtained from the same LSFM datasets, and ultimately to develop an integrated and unbiased quantitative database of identified cellular populations in the human neocortex. This parallel validation approach, performed on separate specimens prepared in thinner materials (50 μm-thick microtome sections of the entire area 44/45) and immunostained in series for each marker of interest, provides layer-specific estimates of cellular features, such as total numbers, local densities, somatic volumes, and spatial distribution of various neuronal populations identified by specific markers including cytoskeletal proteins, calcium-binding proteins, and neuropeptides, as well as generic cellular markers such as NeuN and the Nissl stain. These evaluations serve as a gold-standard for the project. Total neuronal number estimates based on Nissl-stained series of sections from the entire Broca’s area yielded values averaging 35 million neurons in layer 3 and 22 million in layer 5, which is consistent with data obtained independently from prefrontal and temporal regions^67–69^. Preliminary assessment of a subpopulation of large pyramidal neurons identified by non-phosphorylated neurofilament protein (NPNFP) shows that these neurons account for about 30% of total neurons in both layers 3 and 5 of the Broca’s area (**Supplementary figure 3d**), in agreement with previous characterizations of this subset of projection neurons^62,68^. Also, individual Nissl-stained pyramidal cell volumes averaged 2,600 μm^3^ in layer 5 and 2,400 μm^3^ in layer 3, the NPNFP-immunoreactive cells being the largest ones with perikaryal volumes up to 6,000 μm^3^. These preliminary observations are in agreement with previous characterizations of this subset of projection neurons^62,68^.

### 2.6 Multimodal registration for MRI-referenced cell census

A defining feature of this project is processing the same human brain sample through each of the imaging techniques and quantitative tools described above. This enables us to place the cellular anatomical information within a cortical coordinate system. Cross-modality registration of cytoarchitectural properties is complicated due to distortions introduced by histological imaging techniques. While registration is facilitated by leveraging the fact that OCT is performed on the tissue block before sectioning, slices imaged with LSFM show significant distortions when compared to the OCT data. Hence we have developed a non-linear registration method that uses the segmentation of common features visible across all imaging modalities, such as blood vessels, to overcome this challenge. Vessels that have a diameter larger than 150 μm were manually segmented in MRI data and in each LSFM slice, and the Frangi filter^70^ was used to segment vessel-like structures in OCT data (**Supplementary figure 4**). The resulting labels were used in a composite objective function that optimizes intensity similarity and label overlap to drive registration between MRI and OCT on one hand, and LSFM and OCT on the other hand.

We also demonstrate instantiation of microscopic cellular and stereologic information from distorted LSFM histological images within the MRI volume with OCT data serving as a critical intermediary modality with mesoscopic resolution and minimal distortion (**Figure 6** and **Supplementary Movie 3 and 4**). To the best of our knowledge, this is the first time that stereological annotations performed at the microscopic scale are mapped back into the space of the intact brain.

**Figure 6:**
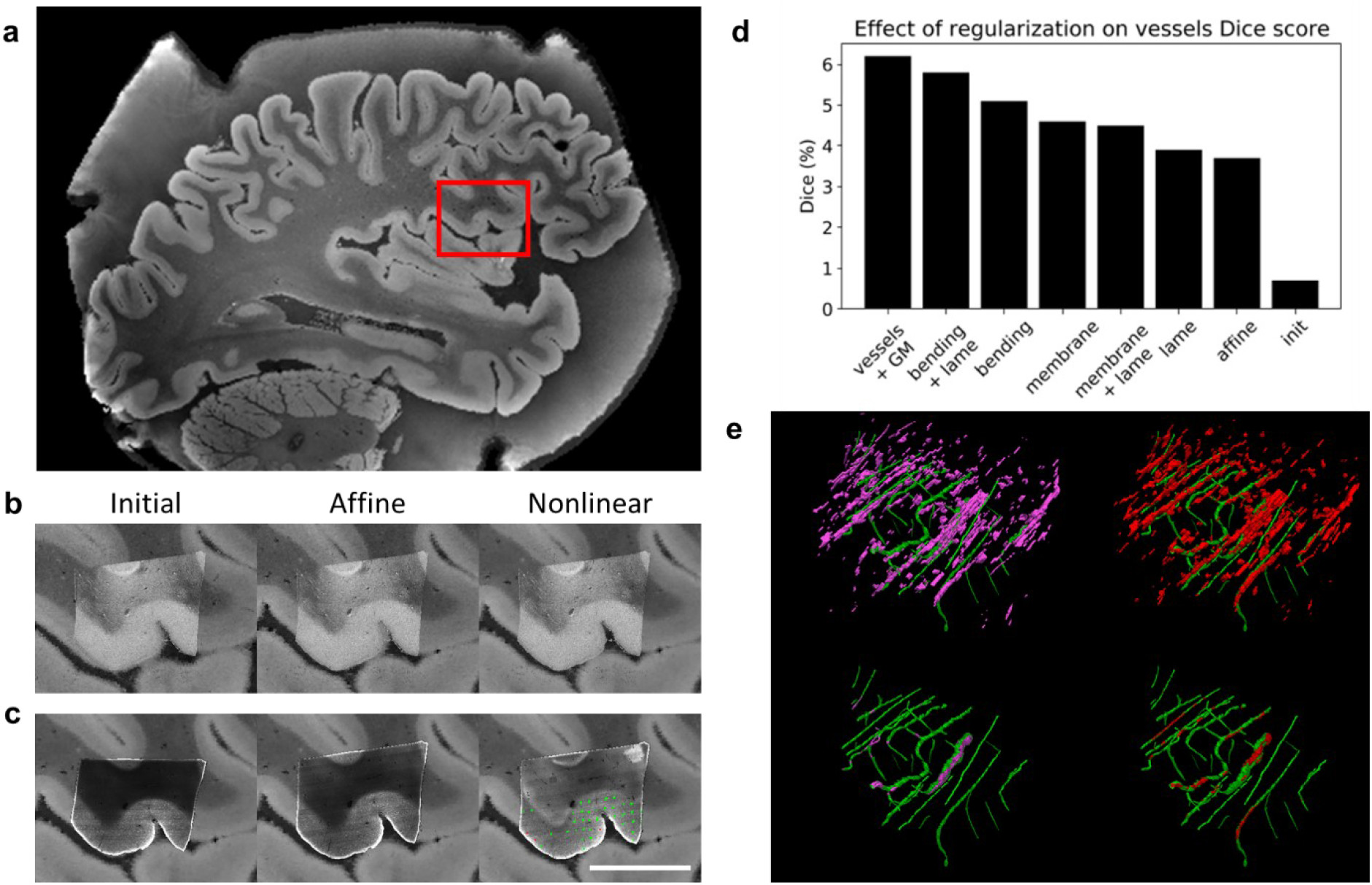
MRI, OCT, LSFM and stereology warped into a single coordinate space. Although the imaging data can be mapped into any space, we chose to display the alignment in OCT space, because it preserves the sectioning axis, allowing all modalities to be visible with minimal distortions. a) Panel showing the whole-hemisphere MRI image with a red box around area 44/45. b) Images showing the MRI deformed to OCT space after initialization, automated affine, and automated nonlinear registration. c) Images showing a slice of LSFM deformed to OCT space after initialization, automated affine, and automated nonlinear registration. The nonlinear registration panel further shows NeuN (green) and CR (red) counting coordinates obtained by stereology and warped to OCT space. d) Graph showing the effect of nonlinear regularization on vessels Dice score for MRI/OCT registration. *init*: initial alignment; *affine*: similitude with 7 degrees of freedom; *membrane*: penalty on first derivatives; *bending*: penalty on second derivatives; *lame*: penalty on zooms and shears, *vessels+GM*: bending+lame with the addition of a vessel- and cortex-based objective function). Scale bar = 1 cm e) Images showing the vessels registration in MRI and OCT with and without a vessel-specific loss. The vessels manually segmented in the OCT volume (green) are shown alongside vessels manually segmented in the MRI volume and warped to the OCT volume using either a purely intensity-based objective function (pink) or a composite intensity- and vessel-based cost function (red).

All deformations are fully invertible and allow any one modality to be warped to the space of any other modality. In addition to vasculature, the boundary between the cortical infra- and supragranular layers was segmented in MRI data and in a subset of LSFM slices to measure registration accuracy. The resulting transforms were used to warp the MRI segments to each manually segmented LSFM section (**Figure 7**), where the 95th (559 ±200 μm) and 75th (310 ±96 μm) percentiles of the minimum distance from the LSFM boundary to the MRI boundary were computed. It should be noted that these distances were computed in 2D sections and are therefore upper bounds on the 3D distances, and that 75% of boundary points have less than two voxels errors in the 150-μm isotropic MRI space.

**Figure 7:**
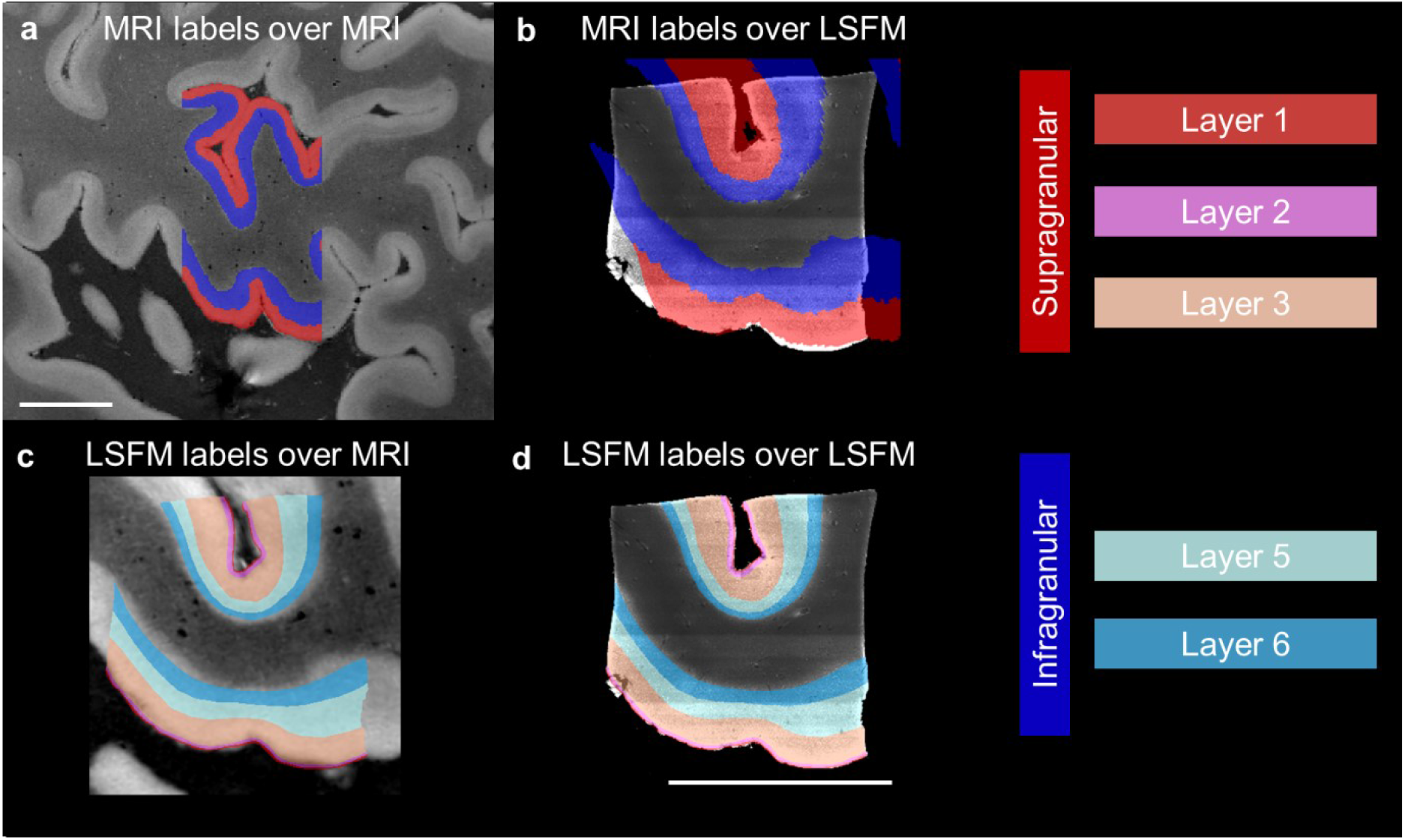
Overlap of Infragranular and supragranular laminar labels derived from MRI and LSFM after coregistration. a) MRI overlaid with the manually labeled infra- and supragranular layers. b) LSFM overlaid with the same MRI-derived labels after registration. c) MRI warped in LSFM space overlaid with LSFM-derived laminar labels. d) LSFM overlaid with manually labeled cortical layers. There is no internal granular layer 4 in Broca’s area. Scale bars = 1cm.

## Discussion

The pipeline presented here demonstrates the possibility of performing a cell census of the human brain, at micrometer resolution, by combining different techniques that overcome the inherent limitations associated with any single modality. To achieve this goal, we implemented new imaging and image processing approaches to correlate the four modalities involved: MRI, OCT, LSFM, and stereology. By first establishing a reference coordinate system for the cellular atlas, through volumetric MRI, we greatly expand the utility of the cellular atlas and provide a basis for *in vivo* inferences with our analysis. Existing 3D models of the human brain with cellular resolution such as the Big Brain project^17^ include sectioning distortions inherent to histology and do not permit through-plane tracing of features of interest across slices such as connecting axons, vasculature and laminar surfaces. However, the ability to build undistorted laminar models of the human cerebral cortex is a critical component of any cellular atlas. In the last decade, the combination of tissue clearing and high-throughput microscopy techniques, such as LSFM, have paved the way for the investigation of brain anatomy in 3D^71,72^ with subcellular resolution. However, clearing methods and physical sectioning of the human brain sample introduce tissue distortions that complicate the coregistration between MRI reference data and fluorescence reconstructions. Attempts to perform MRI on cleared samples showed a contrast loss that prevents visualization of tissue microstructure^73^, suggesting that MRI should be performed before tissue clearing. To facilitate the alignment between MRI and LSFM reconstruction, we decided to use OCT as an intermediate method with cellular resolution to enable registration of the distorted fluorescence images of cleared volumes to the MRI-based atlas coordinate system. We demonstrated for the first time accurate alignment of OCT and whole-hemisphere MRI of the brain at the vasculature level. As OCT data is acquired before slicing, it provides a critical, minimally distorted, intermediate reference between the MRI and LSFM.

Another important aspect of our pipeline is the choice of using immunofluorescence for specific identification of cellular markers both for LSFM imaging and stereological analysis. The precise immunoreactivity in neurons for neurochemical markers allows the definition of region-specific staining patterns, and generally results in a better definition of the cortical areas compared to Nissl or myelin preparations. In this context, markers like neurofilament proteins and calcium-binding proteins are reliable cellular markers for the mammalian neocortex that identify distinct neuronal populations exhibiting recognizable patterns of regional and laminar specialization^35,56–58,60–64,74^. For example, NPNFP is present within a relatively small subset of large pyramidal neurons in layers 3, 5, and 6 that display a highly specific regional distribution^62,74–77^. Similarly, calcium-binding proteins and many neuropeptides are expressed in morphologically non-overlapping subgroups of GABAergic interneurons that are further classified by their electrophysiologic properties and local connectivity^78,79^. Interestingly, while these markers define generic classes of neurons, they also identify neuronal groups known to be differentially affected in many neuropsychiatric conditions. For example, NPNFP-enriched neurons are particularly affected in the course of Alzheimer’s disease while GABAergic interneurons are spared^68,80^, whereas in other conditions such as schizophrenia or autism spectrum disorder the same neurons present different vulnerability profiles^81–84^.

Furthermore, this approach can be integrated with myeloarchitecture data and cerebral microvasculature distribution to enable the determination of patterns of molecular cytoarchitecture and connectivity on multiple scales, from single layers to columnar domains of cortex within a cortical region identified based on specific cytoarchitectural characteristics. To this end, we integrated analytical neurostereology techniques with layer-specific resolution within the pipeline^65,66^. These approaches are rigorous and validated, and have been extensively applied to the study of the human brain^68,69,85–89^. The analysis of multiple fluorescence channels on the same counting frame location is also possible^68^. Such morphometric parameters can be used together with a combinatorial expression profiling analysis of cell classes to provide a comprehensive morphomolecular characterization of the cortical cellular typology^90^. A fundamental aspect of stereologic approaches is the accurate delineation of the region of interest. For Broca’s area, we established reliable cyto- and chemoarchitectural criteria based on published studies^89,91,92^. Finally, stereology also provides a gold-standard against which automated approaches, such as machine learning-based methods to quantify cell types, can be verified. A final aspect of the proposed pipeline is the big data management that was optimized to allow the combination of the different techniques. Indeed, special attention was devoted to the choice of the nomenclature, reference data type format, compression, and post-processing methods to achieve a standardized procedure for data storage and sharing, replicable upon multiple samples.

Several challenges remain to be addressed further to improve the current pipeline. The 3D stereological analysis employed in this study to obtain a reliable counting of cells is highly time-consuming. To obtain the census of the neurons in the entire acquired volumes, the raw images will be automatically analyzed. In particular, we are exploring the possibility of employing a convolutional neural network for pixel classification, previously proposed to analyze two-photon fluorescence microscopy images^93^ and automatic cell detection obtained on mouse brain reconstruction with LSFM^55,94,95^. The approaches will be used to count the different neuronal populations stained in the samples automatically, providing a comprehensive characterization of the whole imaged volume. Moreover, to expand the molecular phenotyping of the tissue, multi-round staining on cleared sample^55,90,96^ will be implemented to characterize human brain cytoarchitecture in greater details. In this context, a quantitative database of morphofunctional neuronal types in identified cortical regions represents a crucial normative resource for the study of cellular changes in brain disorders. The consideration of differential cortical vulnerability in brain diseases can also be used for targeting key cortical domains to carry out future analyses. Also, to speed up the alignment process between the different techniques, eliminating the need to use manual landmarks as performed in this study, microstructure-informed automatic non-linear registration tools are needed. In this regard, we will enhance this pipeline through optimizing an automated vascular segmentation approach^97^. The reference labels will serve as examples to train segmentation models that will be used on future samples, bypassing manual annotation towards the creation of a scalable human brain cellular atlas.

The successful realization of this novel acquisition and analysis pipeline ensures reliable replicability across laboratories, representing a substantial scientific step forward in terms of rigor and reproducibility. We believe that the application of our multimodal approach will provide a deeper understanding of the human brain architecture across resolution levels. Our imaging technology pipeline will ultimately enable the automated reconstruction of undistorted 3D microscopic models of not just a brain area, but of an entire human brain, permitting the assessment of intra- and inter-subject variability. In turn, this will allow investigations of the presence and spread of pathologic cellular alterations observed in neuropsychiatric illnesses in which selective vulnerability of different cell types occur in disease-specific distribution patterns.

## Methods

### Human Brain sample collection

The samples used in this project were obtained from control subjects who died of natural causes with no clinical diagnoses or neuropathology. The brain hemisphere imaged for this paper was from a 79 year-old male. A standard fixation protocol was used in which brain was fixed in 10% formalin for a minimum of 90 days. The sample was packed in a 2% buffered paraformaldehyde solution for MRI scanning. After OCT imaging, tissue slices were stored in phosphate-buffered saline. Human brain tissue samples were procured from the Department of Neuropathology at the Massachusetts General Hospital (MGH) Autopsy Service (Boston, USA). Written consent was obtained from healthy participants prior to death, following IRB-approved tissue collection protocols from Partners Institutional Biosafety Committee (PIBC, protocol 2003P001937).

### MRI imaging

*Ex vivo* MRI is performed on the whole human hemisphere using multiple flip angles of a multi-echo fast low-angle shot (ME FLASH or MEF) sequence at 150-μm isotropic resolution, on a 7 T Siemens MR scanner. Specific scan parameters were: TR = 34 ms, TE = 5.65, 11.95, 18.25 and 24.55 ms, respectively, field of view (FOV) = 192 x 81.3 mm, and slice thickness = 150 μm. A novel acquisition and optimization framework was developed to enhance signal-to-noise ratio (SNR) and correct geometric and intensity distortions without additional high-resolution scans. In the presence of B0 inhomogeneity, the odd and even echoes of the MEF are distorted in opposite directions because they are acquired with opposite polarity readout gradients^49,98^. To remove these distortions, we collected a 2D-encoded B0 fieldmap, which estimates the amount of inhomogeneity at every voxel and in turn gives us a measure of displacement at every voxel of the 150-μm isotropic MEF. We acquired a standard gradient echo fieldmap consisting of two gradient echoes with TR = 5 s, FOV = 192 x 144 mm, matrix size = 160 x 120, and slice thickness = 1.2 mm. Geometric distortions are then corrected using a group sparsity-based edge preserving intensity correction algorithm that uses the fieldmap and all the FLASH images jointly to perform the correction^48^. Additionally, B1+ variation was estimated by acquiring multiple single echo FLASH sequences with short TE (2.7 ms), long TR (5 s), flip angles varying from 20° to 340°, FOV = 192 x 156 mm, matrix = 96 x 78, and slice thickness = 2 mm. We fit the frequency of the sinusoid at each voxel to estimate the multiplicative bias in our flip angle measurements. The estimated flip angle bias map was then used to correct the flip angle value at each voxel location. We fit T1 tissue parameter using a dictionary look up procedure and the corrected flip angle map. We synthesized new FLASH MRI scans with the estimated T1 using the MRI physics forward model for FLASH MRI contrast to remove variations caused by spatially non-uniform B1+ field^49,50^.

The whole hemisphere was imaged using a custom-built 31-channel phased array coil^21^ whose intrinsic sensitivity profile can cause non-uniform intensity. The lack of a body coil with a uniform receive sensitivity profile on the 7 T scanner makes acquisition-based receive bias estimation non-applicable to our scenario. We use a model-based B1- bias correction method that jointly segments tissue type at each voxel and estimates the intensity bias using a likelihood-based cost function. The method assumes that the voxels classified under the same tissue class will have the same FLASH intensity value^51^.

### Serial sectioning OCT imaging

A custom-built polarization-sensitive OCT (PSOCT) system was used to acquire the volumetric data, however, only the intensity signal was analyzed as the polarization part was beyond the scope of this study. Therefore, we use the terminology of OCT instead of PSOCT in the results. The system was built based on a previously reported setup^99^ with a schematic as shown in **Figure 3a**. Specifically, a swept light source (AxsunTech) was used in the OCT system which has an A-line rate of 100 kHz, a center wavelength of 1310 nm and a bandwidth of 135 nm. The axial resolution was estimated to be 5.6 μm in tissue (with a refractive index of 1.4). The sample arm consists of a pair of XY scanning mirrors, a 4× telescope, and a 4× air objective (Olympus, UPLFLN4x, NA 0.13) to obtain a lateral resolution of 6 μm. The interference fringe from the sample and the reference arms was collected by a balanced detector. The post-objective power was measured to be 3.7 mW, achieving a 95 dB SNR. For this study, we used a 3×3 mm^2^ FOV with a 3-μm lateral step size and 10% overlap between tiles, with each tile takes about 16 s. For large human brain blocks, the dimension of the embedded samples are usually a few centimeters on each dimension, which is over ten times greater than that of the FOV of a single image tile. Therefore, the whole sample surface was divided into a grid of views and the images from all views were stitched together to form a full surface. Motorized xyz stages (x and y stages: LTS150, Thorlabs; z stage: MLJ150, Thorlabs) were incorporated to translate the samples under the OCT scanning head to all the imaging locations. The maximum travel distance for x, y and z stages was 150 mm, 150 mm and 50 mm with correspondent one-direction moving accuracy of 2 μm, 2 μm and 10 μm. A customized vibratome slicer was mounted adjacent to the OCT imaging head to cut off a superficial slice of the tissue block upon completion of the scanning of the sample surface. A 2.5-in custom sapphire blade (DDK) was vibrated at 50 Hz and 1.2 mm peak-to-peak amplitude while slicing, with 0.1 mm/s stage feed rate. Custom software written in LabVIEW was used to control OCT imaging and vibratome slicing.

The data acquisition computer was a high-performance local computer with a four-core processor, 64 GB memory, a high-speed digitizer (ATS9350, Alazar), a GPU (RTX4000, NVIDIA) and a 10 GB/s high-speed Ethernet card. Using the k-clock from the light source, the signal was digitized in even-k space. The GPU fetched the spectral domain data and performed real-space reconstruction on the fly, which included dispersion compensation^100^, Fourier transform for depth-profile creation and rough trimming in depth. The reconstructed data was then saved to a local storage server with 28 TB space through the 10 GB/s Ethernet. Compared with the 0.2 GB/s data acquisition rate, the Ethernet transfer rate was much faster and helped avoid any data loss. For high-speed post processing, data saved in local server was automatically uploaded to Boston University Shared Computing Cluster (SCC), a high-performance computing resource located at the site of Massachusetts Green High Performance Computing Center at Holyoke, Massachusetts. Upon completion of the experiment, a parallelized post-processing script was executed on SCC which included distortion correction, volume stitching and various features extraction. We utilized both ImageJ plug-ins^101,102^ and customized functions to correct distortion that is introduced during OCT acquisition, such as shading effect and gird distortion. To stitch the OCT volume in 3D, we first stitched tiles in 2D using the average intensity projection. After the coordination for each tile is obtained, we linearly blended them in x and y and then stacked them in z. Once the OCT volume is reconstructed, various features, such as vessels, axonal bundles and cortical laminar structure can be extracted using feature enhancement algorithms.

### Tissue clearing and labelling for LSFM imaging

Brodmann’s areas 44/45 brain slices were treated with SHORT^55^ a modified version of the SWITCH/TDE tissue transformation protocol from Costantini et al.^93^, that combines the SWITCH technique^90^ with the TDE clearing method^103^. Briefly, each slice was incubated in a SWITCH-off solution, consisting of 50% phosphate-buffered saline (PBS) titrated to pH 3 using HCl, 25% 0.1 M HCl, 25% 0.1 M potassium hydrogen phthalate, and 4% glutaraldehyde. After 24 h, the solution was replaced with PBS pH 7.4 with 1% glutaraldehyde. Finally, the samples were incubated in the clearing solution for 2-4 days at 55 °C. The presence of lipofuscin in the cellular soma^104^ and the free-aldehyde double-bounds introduce high autofluorescence signals^105^. To decrease such spurious and non-specific signals, the specimens were treated with hydrogen peroxide (30% v/v) for 1 h at room temperature (RT). After several washes in PBS, antigen retrieval was performed using pre-heat tris-EDTA buffer (10 mM Tris base (v/v), 1 mM EDTA solution (w/v), 0.05% Tween 20 (v/v), pH 9) for 10 min at 95 °C. To perform the multicolour staining, after 3 washing steps with deionized water (DI) and rebalancing in PBS for 1 h, the samples were incubated with primary antibodies against NeuN (Merck ABN91 chicken) and calretinin (CR); Proteintech 12278-1-AP rabbit) at 37 °C for 4 days in PBS +0.1% Triton (PBST). Dilutions for the anti-NeuN and -CR antibodies were 1:100 and 1:200 respectively). Following several washes in PBST, the samples were incubated for four days with the secondary antibodies conjugated with different Alexa Fluor dyes (dilution of 1:200). The slices were rendered transparent by soaking the samples in increasing solutions of 20%, 40% and 68% (vol/vol) of 2,2’-thiodiethanol in PBS, each added with propidium iodide (dilution of 1:100) for 1-2 days at RT with gentle shaking. Samples were mounted in a sandwich-like configuration between a 250 μm thin quartz coverslip (for refractive index matching at a refractive index of 1.46) and a standard glass sample holder, with a 500 μm-thick steel spacer in between^55^. Glycerol (91%) in distilled water was used outside the sandwich for the LSFM objectives immersion. This step permitted us to achieve high penetration depth and to avoid any optical aberration by matching the refractive index of the brain samples.

### LSFM imaging

In our custom-made inverted light-sheet fluorescence microscope^54^, two identical objectives were inclined at 90° from each other and were spaced such that their fields-of-view (FOV) are orthogonal and overlap in the center. They alternately play excitation and detection roles. The objectives were a pair of LaVision Biotec LVMI-Fluor 12x PLAN with 12x magnification, NA 0.53, WD 8.5-11 mm, spherically and chromatically corrected in the visible range, with a correction collar for refractive index matching with the immersion solution. They were inclined at 45° relative to the sample holder plane to allow for the largest possible lateral sample size while not interfering with its extension within the plane. These objectives were carefully chosen to maximize the optical resolution (1.1 μm lateral and 3.7 μm axial) and field of view (1.1×1.1 mm^2^), while respecting the geometrical constraints and allowing immersion in any refractive index matching media. The microscope was equipped with four laser sources (Cobolt MLD 405 nm/100 mW, MLD 488 nm/60 mW, DPL 561 nm/100 mW, MLD 638 nm/180 mW), each emitting a Gaussian beam that had its width adjusted by a dedicated telescope, before combining all of them through a set of three dichroic mirrors. This combined beam was split by a 50-50% beam splitter in two equal parts that were conveyed into the two identical excitation pathways of the light-sheet microscope. In each pathway the beam was modulated in intensity, timing and transmitted wavelength by an acousto-optical tunable filter (AOTF, AAOptoelectronic AOTFnC-400.650-TN) and then was scanned by a galvo mirror (Cambridge Technology 6220H), in order to realize the digitally scanned light sheet planar illumination^106,107^. A scanning lens (Edmund Optics #45-353, fL=100 mm, achromat), placed after the galvo mirror, converted the angular deflection into a lateral displacement of the incident light. The beam was then directed by the excitation tube lens (Edmund Optics #45-179, fL=200 mm, achromat) to the objective’s pupil, through which it sequentially illuminated neighboring lines within a single plane of the sample. Each objective was held on a motorized stage (PI L-509.14AD00) to adjust its focal plane position. The sample was held in a custom quartz sample holder inserted into a plastic tray filled with refractive index matching medium. The sample was positioned using a 3-axes motorized stage system (two PI M-531.DDG and a PI L-310.2ASD for a motion range within 30×30×2.5 cm^3^ with submicrometric repeatability) and was imaged by translating it along the horizontal direction. The image velocity through the volume was 47 frames/s, corresponding to a volumetric rate of 0.5 cm^3^/hour. The fluorescence emitted by the sample was collected by the other objective lens, then was separated from the reflected laser excitation light by a multi-band dichroic beam splitter (Semrock Di03-R405/488/561/635-t3-55×75), before being directed by the detection tube lens (Edmund Optics #45-179, fL=200 mm, achromat) on a sCMOS camera (Hamamatsu OrcaFlash4.0 v3). Each camera operated in confocal detection mode by having the rolling shutter sweep in synchrony with the galvo scan of the digital light sheet^95,108–112^. Five sets of band-pass filters were mounted in front of each camera on a motorized filter wheel (Thorlabs FW102C) to image selectively the differently labeled cells or structures within the tissue sample.

The operation of the microscope hardware was controlled by a workstation running a custom multi-threaded software, developed in C++ using Qt with a flexible and modular architecture and composed now of approximately 9000 lines of code. Our software ensured hardware synchronization and triggering by using a National Instruments PCIe-6363 card and controlled the automatic image acquisition from the two sCMOS cameras in confocal detection mode, with a sustained data rate of 800 MB/s at 47 fps and storage on a 16 TB SSD RAID.

The acquisition procedure for any sample started from determining its edges. Each image stack was acquired by moving the sample along the x axis through the fixed FOVs of the two objectives, then the sample was shifted by 1 mm along the y axis and the next stack was acquired. This sequence continued until the whole volume had been acquired. Contiguous stacks had an overlap of 100 μm that allowed to fuse them in post-processing to form the whole volume. Appropriate metadata was saved jointly with the acquired stacks.

The two identical optical pathways of the LSFM alternately served as excitation and detection arms, with a time-delay of a half frame that was introduced between the two roles to avoid exposing the active rolling-shutter rows on the acquiring camera to stray light coming from the illumination beam on the same side. The two AOTF allowed to shutter each illumination pathway independently to avoid introducing stray light and, furthermore, enabled to select which laser wavelengths and intensities were impinging on the sample.

### LSFM data management

To visualize the reconstruction of an entire slice, a post-processing pipeline was applied to the data. First, as the objectives acquire images of the moving sample at 45° relative to the slide plane, an affine transform was applied to compensate for the motion and the 45° rotation thus bringing the acquired volume back to the sample’s coordinate system. The affine transform also performs a spatial down-sampling to 3.3-μm^3^ isotropic resolution. Then a custom-made stitching software, ZetaStitcher (G. Mazzamuto, “ZetaStitcher: a software tool for high-resolution volumetric stitching” https://github.com/lens-biophotonics/ZetaStitcher), allowed us to fuse the contiguous stacks, using the overlapping regions, to obtain a representation of the whole sample. Only for visualization, an illumination intensity homogenization algorithm was applied to the stitched volume or to single slices to compensate variations in the laser beam power across the field of view and among stacks. For each fluorescence band, the observed intensity along the light propagation axis was averaged, attaining a smooth intensity profile. By dividing each image for this reference, illumination intensity artefacts occurring across the transversal sample extension were mitigated. To store and share the information, data was compressed using the JPEG2000 lossy approach that yields a 1:20 compression. The data analysis pipeline was written in Python. The 561 nm wavelength 3D reconstructions were used to visualize and segment the blood vessels to perform the alignment of the three modalities as described in the section below.

### Stereology

Stereologic analysis was performed on each cleared slices of Broca’s area, 500 μm-thick, imaged at a 3×3×3 μm pixels dimensions, and immunostained for NeuN and calretinin, using the MBF Bioscience Stereo Investigator Cleared Tissue software (version 2020.1.1) with an Optical Fractionator design^65,66^. The counting frame size was 150 x 150 μm, the grid size was 800 x 800 μm, and the disector height was 15 μm for all sections of tissue examined generating >600 sampling sites. Layers 3 and 5 were outlined and their boundaries were used to estimate laminar surface areas and volume. There were 10 virtual 49.5 μm-thick sections for the tissue sample and layers 3 and 5 were contoured at the widest part of each sub-slab, with a 400% zoom. Markers were placed at the top of each sampled cell, as it came into focus within the depth of the disector.

### Image registration

Intermodality deformations were modelled using a combination of affine and nonlinear transformations. The affine model was restricted to a similarity transform (i.e., combinations of translations, rotations and isotropic scaling) and encoded in the Lie algebra of the corresponding conformal Euclidean group^113^. Non-linear transformations were modelled by stationary velocity fields and exponentiated using a scaling and squaring algorithm^114^, which ensures – under mild smoothness conditions – that the resulting transformations are invertible diffeomorphisms. The parameters of the deformations were optimized by minimizing a combination of losses on intensity images (normalized mutual information^115^) and manually segmented landmarks (soft Dice^116^). Stationary velocity fields were regularized with a combination of penalties on their bending and linear elastic energies^117^. When registering a whole volume (in this case, MRI) with a sub-block (e.g., OCT), the larger volume was always deformed to the space of the smaller block, where the objective function was computed. Because the transformation model is diffeomorphic, the resulting transforms could nevertheless be inverted and used to warp the block back to the space of the larger volume. The same transforms were also used to warp stereology coordinates extracted from the MBF software. The registration model was implemented in PyTorch^118^, leveraging automatic differentiation, and jointly optimized with Adam^119^. Parameters were progressively optimized in a coarse to fine fashion (rigid, then affine, then affine and nonlinear with a progressively finer grid), and the learning rate was divided by 10 when the objective function reached a plateau, until convergence. Specifically, for MRI to OCT registration, the OCT volume was downsampled to 100 μm, and normalized mutual information (NMI) was computed within patches of 150×150×150 voxels. The objective function combined NMI (weight 1), Soft Dice between grey matter segments (weight 1), Soft Dice between vessels segments (weight 0.1) and regularization (bending energy: 4, divergence: 1, shears: 4). For OCT to LSFM registration, the OCT volume was downsampled to 50/100/200 μm and the LSFM volumes were downsampled to 20/40/60 μm. The objective function was computed jointly at all resolutions and combined NMI (weight: 2), Soft Dice between vessels segments (weight: 0.1) and regularization (bending energy: 0.4, divergence: 0.1, shears: 0.4). The effect of different components of the SVF regularization was investigated in an ablation study. Dice scores between manually segmented vessels in the MRI and OCT volumes were computed in the initial position (*init*), after affine registration (*affine*) and after nonlinear registration with different composite types of regularization: the membrane energy (*membrane*) penalizes first derivatives, the bending energy (*bending*) penalizes second derivatives and the linear-elastic energy (*lame*) penalizes local shears and zooms. The best Dice score was obtained with a combination of bending and linear-elastic penalties.

## Acknowledgements

We thank Jack Glaser and Julie McMullen (MBF Bioscience) for all their help during implementation of the Stereo Investigator Cleared Tissue software in our application. We express our gratitude to the donor involved in the body donation program of the Massachusetts General Hospital who made this study possible by generously donating his body to science. Support for this research was provided in part by the BRAIN Initiative Cell Census Network grant U01 MH117023, the National Institute for Biomedical Imaging and Bioengineering (P41 EB015896, R01 EB023281, R01 EB006758, R21E B018907, R01 EB019956, P41 EB030006, R00 EB023993), the National Institute on Aging (R56 AG064027, R01 AG064027, R01 AG008122, R01 AG016495), the National Institute of Mental Health (R01 MH123195, R01 MH121885, RF1 MH123195), the National Institute for Neurological Disorders and Stroke (R01 NS0525851, R21 NS072652, R01 NS070963, R01 NS083534, U01 NS086625, U24 NS10059103, R01 NS105820), Eunice Kennedy Shriver National Institute of Child Health and Human Development (R21HD106038), Chan-Zuckerberg Initiative DAF an advised fund of Silicon Valley Community Foundation grant number 2019-198101, and was made possible by the resources provided by Shared Instrumentation Grants S10 RR023401, S10 RR019307, and S10 RR023043. Additional support was provided by the NIH Blueprint for Neuroscience Research (U01 MH093765), part of the multi-institutional Human Connectome Project. This project has also received funding from European Union’s Horizon 2020 research and innovation Framework Programme under grant agreement No. 654148 (Laserlab-Europe); European Union’s Horizon 2020 Framework Programme for Research and Innovation under the Specific Grant Agreement No. 785907 (Human Brain Project SGA2), No. 945539 (Human Brain Project SGA3) and under the Marie Skłodowska-Curie grant agreement No. 793849 (MesoBrainMicr); Italian Ministry for Education in the framework of Euro-Bioimaging Italian Node (ESFRI research infrastructure); “Fondazione CR Firenze” (private foundation), European Research Council (Starting Grant 677697, project BUNGEE-TOOLS), Alzheimer’s Research UK (Interdisciplinary grant ARUK-IRG2019A-003) and NIH R01 AG070988-01 and RF1 MH123195-01.

The content of this work is solely the responsibility of the authors and does not necessarily represent the official views of the National Institutes of Health and other funding agencies.

## Author Contributions

DV developed the B0 and B1 transmit distortion correction methods and the associated imaging protocols to improve laminar and vessel contrast, optimized the scan protocol to improve overall CNR and SNR of MRI and integrated intensity bias correction method into the processing pipeline. AVDK developed the custom sequences used to acquire high resolution structural MRI data, built the custom data-handling infrastructure and oversaw the development of the MRI scan protocol with BF. DV, RF and LM contributed towards the development of the MRI scan protocol, image acquisition, and analysis. YB contributed towards the development of the multi-modal registration pipeline, building on original work done by JEI and AD. MFr provided the human brain samples for this study. JA consulted on sample storage and treatment for preservation throughout the imaging pipeline, including consulting on histology protocols.

LM, DC, MVe, JN, KN, JM, EP, and KE performed the *ex vivo* imaging and analysis team and are involved in image acquisition, data processing, and developing standards for segmenting vasculature and the infra/supragranular boundary of the cortex. In MRI, OCT and LSFM data. DC, JN, KN, JM, EP, and KE performed manual labeling of these data. Labeling standards built on CM’s original groundwork for segmenting features of interest for the purpose of registering MRI to OCT data. CM also consulted on OCT imaging and analysis for the BU team. LM oversaw the integration of the image and analysis pipeline and data publication for this project. This builds on MFo’s original project oversight and management.

JY, SCha, HW, and IC developed and performed image acquisition and data processing of OCT measurement. JY, SCha, and SChe built the serial sectioning vibratome. JY coordinated the work on OCT sample preparation and transportation.

IC, LP, and MS developed and performed the clearing and staining protocol; VG, LS, and GM built the LSFM hardware and software apparatus, LP and VG performed light-sheet imaging. GM, FMC, VG, and MR contributed to image processing and data analysis. IC coordinated the work on sample preparation, image acquisition and data processing of LSFM measurement. BW, MVa, SR and JL processed tissue for stereology and performed the stereologic analyses. FSP, DAB, BF, and PRH conceived and supervised the study. BF helped design many of the algorithms used to analyze the data. IC, LM, JY, YB, DV, and PRH, wrote the paper with inputs from all authors.

## Data Availability

All the datasets acquired for this study are made available on the DANDI platform at the link: https://gui.dandiarchive.org/#/dandiset/000026^47^.

