## Supplementary Information for "A multimodal imaging and analysis pipeline for creating a cellular census of the human cerebral cortex"

#### Supplementary figures:

**Supplementary figure 1:** a) Orthogonal views of a representative slice stained with CR in blue, PI in green, and NeuN in red (xy plane displays the reconstruction at ≈180µm depth). b) 3D rendering of the slice in a (Dimension 1500 x 1300 x 0,5 mm). c) 3D rendering of the yellow box in b (Dimension = 2 x 2 x 0,5 mm). (b) and (c) images were made using the plugin "3D Viewer" of Fiji.

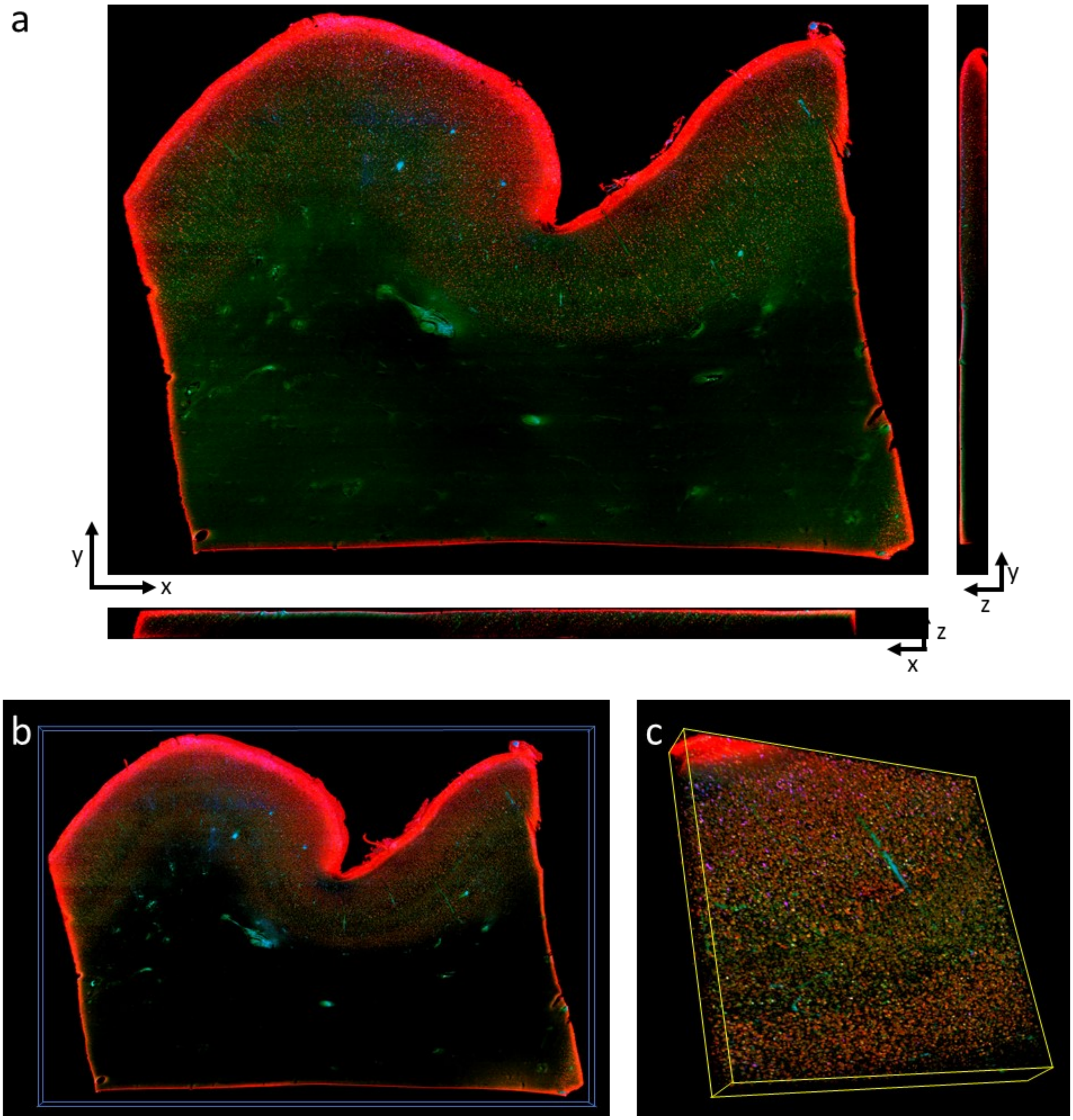

**Supplementary figure 2:** Stereologic quantification of CR$^{+}$ interneurons from LSFM images a) A slab immunostained for CR from the LSFM imaging dataset displaying a systematic-random counting grid employed during stereologic analyses. b) Zoomed-in image displaying cortical layer 3. c) Insets showing the native resolution (3.3 μm isotropic) along the three axes. Grid size on (a) = 800 μm, scale bars on (c) = 200 μm.

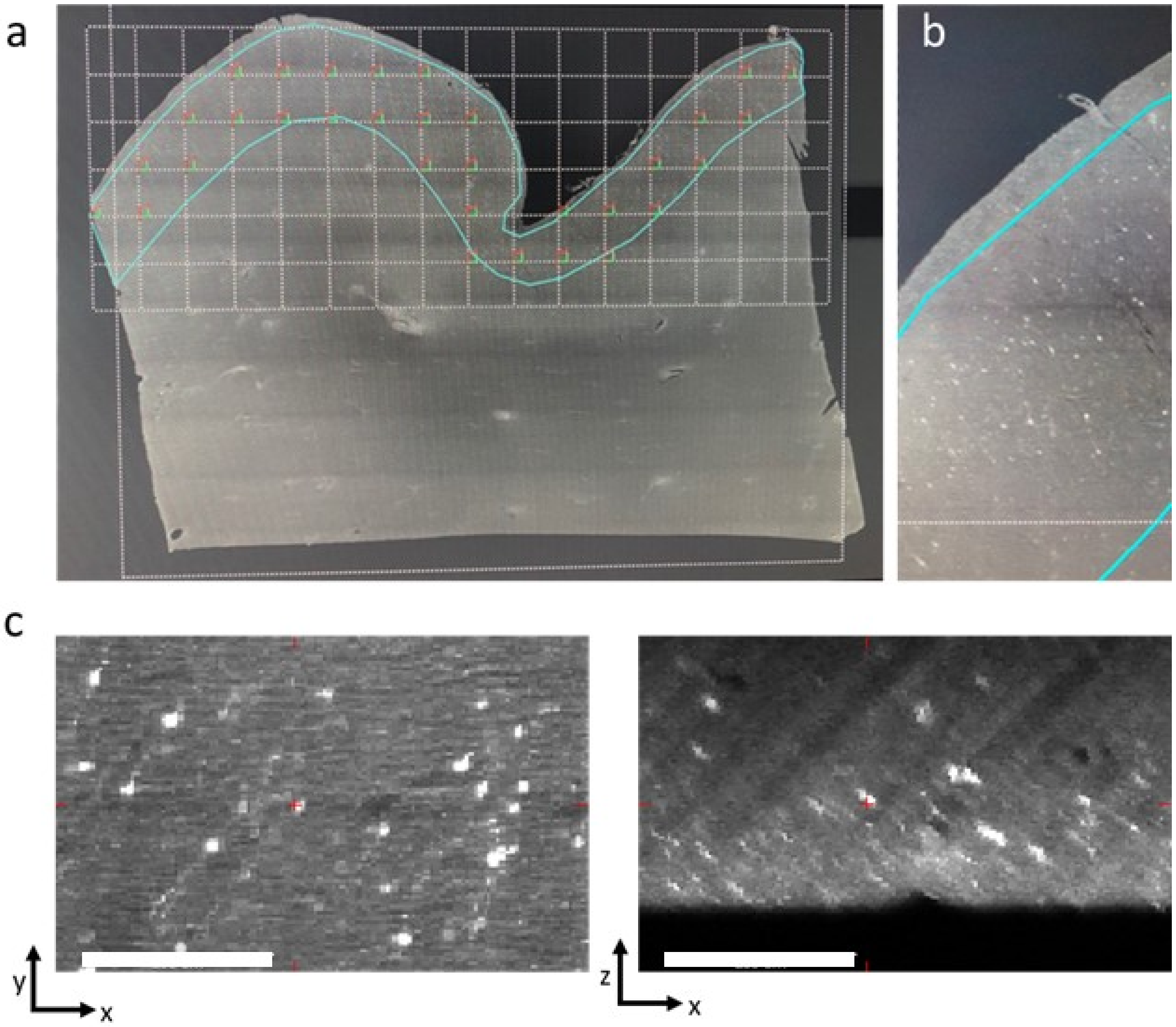

**Supplementary table 1:** 3D stereology counting of calretinin$^{+}$ and NeuN$^{+}$ cells in LSFM images for each virtual slice of the slab from Broca's area evaluated in the study

| **Calretinin** | | **NeuN** | | |
|---|---|---|---|---|
| **Slice #** | **Layer III Estimated Population** | **Slice #** | **Layer III Estimated Population** | **Layer V Estimated Population** |
| 16 | 24499.20 | 16 | 90412.38 | 65681.63 |
| 17 | 11642.40 | 17 | 95556.27 | 74060.80 |
| 18 | 10851.87 | 18 | 111372.80 | 87108.27 |
| 19 | 22903.47 | 19 | 93678.93 | 66926.93 |
| 20 | 20744.53 | 20 | 83729.06 | 53785.60 |
| 21 | 20744.53 | 21 | 94242.13 | 70493.87 |
| 22 | 12108.80 | 22 | 79223.47 | 60731.73 |

**Supplementary figure 3:** Classical stereology evaluation of a thin section. Immunohistochemical detection of microtubule associated protein 2 (MAP2, black) in a 50-µm thick section from Broca's area (a). Counterstaining with cresyl violet enabled detection of cortical layers indicated by their contours as follows: layers 2 and 3 (orange), layer 5 (pink) and layer 6 (cyan). The red markers indicate MAP2-immunoreactive neurons counted in layer 3 and blue markers indicate those counted in layer 5. Panels b and c show examples of immunostaining at higher magnification for MAP2, panel d for non-phosphorylated neurofilament protein and panel e for parvalbumin, respectively. Scale bar on (a) = 2mm, counting frame on (b) and (c) = 100 µm, scale bar on (e) = 25 µm for (d), and 10 µm for (e).

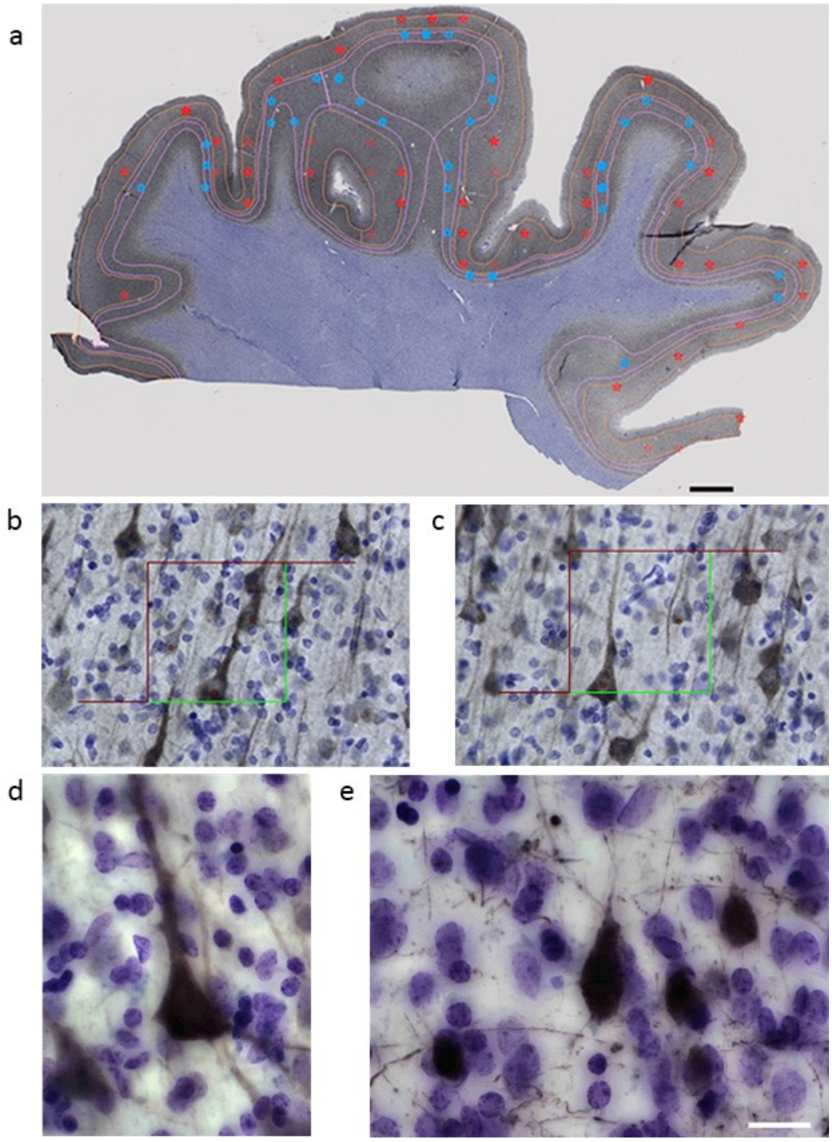


**Classical stereology:**

**Supplementary Method:** All human brains for standard stereology were hemisected at autopsy (postmortem delay 2–12 h) and were obtained from patients without history or symptoms of neurodegenerative or neuropsychiatric disorders at time of death, and the cerebral hemispheres were fixed in 4% paraformaldehyde for up to 3 weeks. The entire inferior frontal gyrus including Broca's area was dissected out, placed in phosphate buffer and then cryoprotected in graded sucrose to 30%. The block was then placed on OCT medium prior to freezing and sectioned on a Leica sliding microtome at a 50-µm thickness. Series of sections were collected for stereology. Following heat-induced antigen retrieval in 10 mM sodium citrate, pH 6.0 with 0.05% Triton X-100, pretreatment with a 3:1 vol:vol mixture of methanol/$H_2O_2$ to prevent endogenous peroxidase reaction and blocking with 5% normal goat serum, series were immunostained, in the example shown in Supplementary Figure 3a-c, with a rabbit polyclonal antibody against MAP2 (Atlas Antibodies HPA008273, 0.1 mg/ml), and incubated overnight at 4°C. Specific labeling was revealed with a biotinylated goat anti-rabbit antibody (1:200; Vector Laboratories BA1000), using the ABC kit (Vector Laboratories), and 3,3'-diaminobenzidine as a chromogen. Immunoreactivity was further enhanced using $NiCl_2$. Other markers were also investigated such as non-phosphorylated neurofilament protein (mouse monoclonal, Biolegend, SMI-32, 1:1000) and calcium-binding proteins (parvalbumin, mouse monoclonal, Millipore Sigma MAB1572, 1:8000) (Supplementary Figures 3d-e and see main text). Materials were then Nissl-counterstained with cresyl violet, which adds a standard definition of laminar cytoarchitecture and permits an assessment of total cell (or neuron) counts and cell individual volumes at the same time a specific population of cells is quantified. For these analyses standard stereologic probes are used under brightfield microscopy using the Stereo Investigator software (v.11 MBF Bioscience) and include the Cavalieri estimator for regional and laminar volumes, optical fractionator for estimates of layer-specific cell numbers and densities, and nucleator for individual cell volume estimates. Inter-rater testing is implemented during pilot studies necessary to establish the parameters to be used throughout the study

**Supplementary figure 4**: Vasculature segmentation: vessels with a radius larger than 100 µm were manually segmented in the MRI volume (a) and in each LSFM slice (c). A semi-automated method based on the Frangi filter was used to segment vessel-like structures in the PSOCT volume (b). Segmented vessels in each modality are also shown in 3D (d: MRI, e: OCT, g: LSFM). Panel f displays LSFM vessels (in green) across all slices after reconstruction by non-linear coregistration with the PSOCT volume, overlaid with PSOCT vessels (in red).

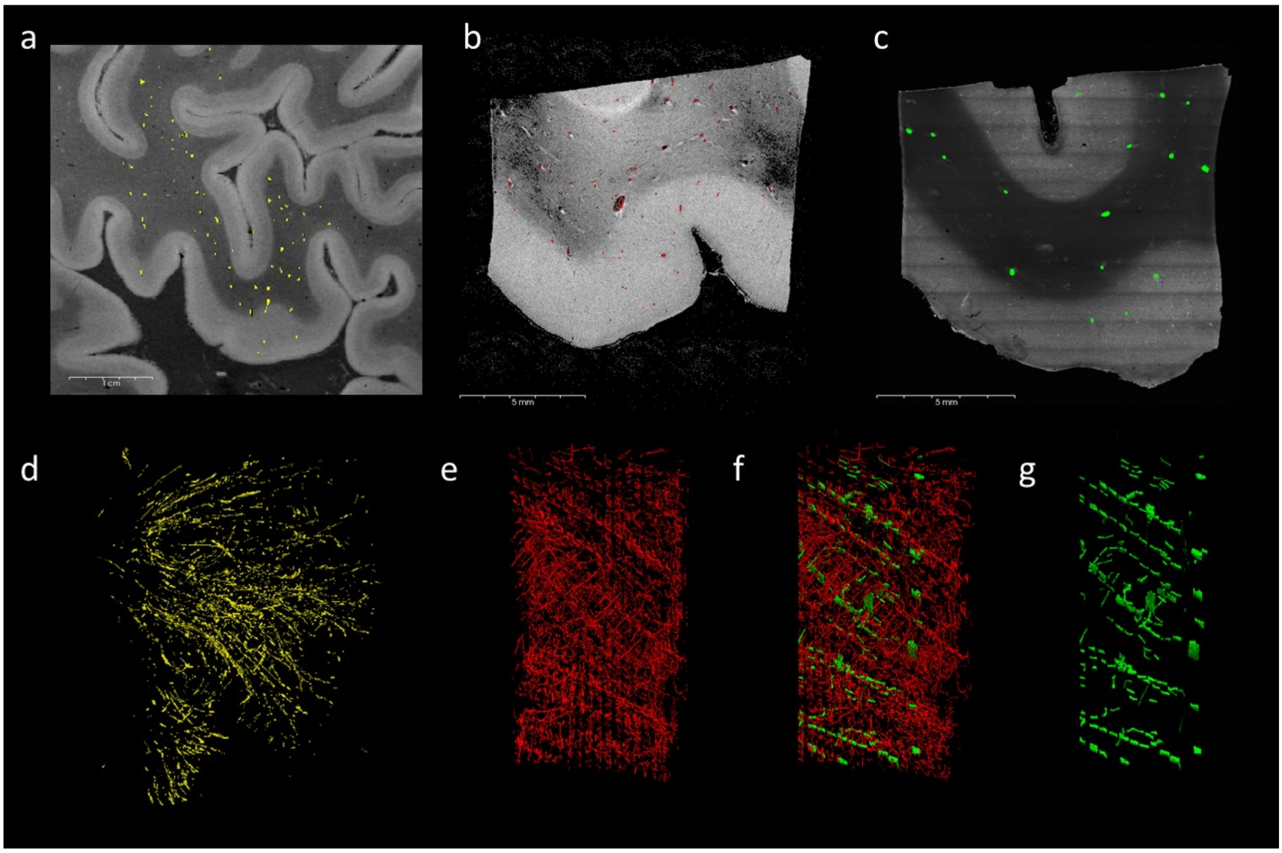


### Supplementary movies:

**Supplementary movie 1**: 3D rendering of the 16 slices acquired with OCT for a total volume of 1.5x1.3x0.8 $cm^3$

**Supplementary movie 2**: Navigation in the 3D rendering of a representative slice acquired with LSFM and stained with anti-Calretinin antibody in blue ($\lambda_{exc}$ =488), Propidium Iodide (nulei) in green ($\lambda_{exc}$ =561), and anti-NeuN antibody in red ($\lambda_{exc}$ =638).

**Supplementary movie 3**: Navigation across sagittal MRI slices and zoom into the Broca's area. Co-registered slices of OCT, LSFM (stained for Propidium iodide, Calretinin and NeuN) and manually segmented cortical layers are displayed in the same space.

**Supplementary movie 4**: Navigation across sagittal MRI slices in Broca's area, with co-registered stereological counts of NeuN-positive cells (green) and Calretinin-positive cells (red) overlaid.
